# Hyperspectral Sensing for High-Throughput Screening of Boron Tolerance in Grapevines

**DOI:** 10.1101/2025.03.21.644478

**Authors:** Yaniv Lupo, Sadikshya Sharma, Jose Munoz, Veronica Nunez, Ana Gaspar, Andrew J. McElrone, Luis Diaz-Garcia

**Affiliations:** Department of Viticulture and Enology, University of California Davis, CA 95616, USA; Crops Pathology and Genetics Research Unit, USDA-ARS, Davis, CA, 95618, USA

## Abstract

Boron (B) is an essential micronutrient for grapevine growth, yet excessive levels can impair photosynthesis, reduce yields, and diminish fruit quality. In this study, we evaluated the potential of hyperspectral radiometry combined with machine learning to identify B-tolerant rootstocks rapidly and cost-effectively. We screened both commercial grapevine rootstocks and wild *Vitis* germplasm under B treatments ranging from 0.5 to 8 ppm, measuring leaf B accumulation, stomatal conductance, photosystem II efficiency, and leaf reflectance (R_380_–R_1100_ nm). Our results revealed substantial genotypic variation in B exclusion, with some genotypes maintaining low leaf B content despite high external concentrations. Classification models (Partial Least Squares Discriminant Analysis and Random Forest classification) outperformed regression models (Partial Least Squares Regression and Random Forest regression) in distinguishing B-excluding genotypes, achieving moderate to high accuracy within just eight days after stress initiation. Vegetation indices such as Normalized Difference Vegetation Index (NDVI), Photochemical Reflectance Index (PRI), Structure Insensitive Pigment Index (SIPI), and Chlorophyll Index (CI) indicated that B stress reduces chlorophyll levels and may induce carotenoid accumulation, suggesting a photosynthetic tolerance mechanism. Although quantitative prediction of leaf B content proved more challenging, simulations showed that even modest prediction accuracies can substantially boost genetic gains if larger populations are screened, and selection intensities are increased. These findings underscore the value of hyperspectral radiometry for high-throughput phenotyping, allowing breeders to rapidly identify and advance B-tolerant rootstocks.

## Introduction

Boron (B) is an essential micronutrient that plays a crucial role in plant growth and development (Brdar-Jokanović, 2020; Brenchley & Warington, 1927; Warington, 1923). In the soil solution, B is primarily available to plants in the form of boric acid (H_3_BO_3_), which is taken up by roots through passive diffusion and active transport via specific transporter proteins (Camacho-Cristóbal et al., 2008). Once absorbed, B is primarily transported through the xylem via the transpiration stream, moving from the roots to the apical parts of the shoots, where it accumulates. While some plant species can also transport B through the phloem to supply non-transpiring organs (Brown & Hu, 1996; Brown & Shelp, 1997; Shelp et al., 1995), it is unclear whether grapevines rely on this pathway (Blevins & Lukaszewski, 1998).

Plants require only minimal amounts of B, with an optimal soil concentration ranging between 0.5 and 2 parts per million (ppm; Botelho et al., 2022). Concentrations exceeding 5 ppm can lead to toxicity in crops (Eaton, 1935; Gunes et al., 2006; Oertli, 1960; Quartacci et al., 2015). Symptoms of B toxicity typically include reduced leaf size, chlorosis, and necrosis at the tips of mature leaves, as B accumulates in leaf tissues (Ozturk et al., 2010; Princi et al., 2016).

High B concentrations are particularly concerning for viticulture in California’s Suisun, Sacramento, and San Joaquin Valleys—regions that collectively produce the majority of the nation’s grapes, including most of the table grapes and raisins consumed in the U.S. (Yau & Ryan, 2008). Excessive soil B in these key agricultural areas can stunt vine growth, reduce yield, and ultimately lower crop value for growers. Given the economic significance of these regions, addressing B toxicity is a grower priority. A deeper understanding of B tolerance mechanisms, the identification of tolerant germplasm, and the development of advanced breeding strategies will be critical to sustaining productivity and fruit quality in grapevine cultivation.

B tolerance is believed to involve the upregulation of B efflux channels that actively transport excess B from roots back into the soil (Reid, 2014). In *Arabidopsis*, elevated B concentrations trigger increased channel activity in root epidermal cells, reducing B accumulation in both roots and shoots (Miwa et al., 2007). Further research suggests that, in rice, B tolerance also involves specific aquaporins, such as *OsPIP1;3*, responsible for B influx, and *OsPIP2;6*, which promotes rapid efflux. The expression of these aquaporins in roots has been shown to increase under elevated B conditions, correlating with improved B tolerance (Mosa et al., 2016). In grapevines, B accumulation increases as soil B concentration and leaf age increase, leading to chlorosis, necrosis, reduced photosynthesis, lower stomatal conductance (g_s_), and stunted shoot growth (Pech et al., 2013; Yermiyahu et al., 2006). These detrimental effects underscore the importance of using B-excluder rootstocks in high-B soils to mitigate toxicity and sustain vine health. Despite its importance, data on B tolerance in grapevines, either commercially available material or wild germplasm, is scarce.

B concentration in leaf samples is typically estimated using either a colorimetric assay or spectrometry. In the colorimetric method, leaf tissues are first digested— usually with acids such as nitric and perchloric acid—to release B into solution. The extract is then combined with an azomethine-H reagent under acidic conditions, forming a colored complex. The intensity of this color, measured spectrophotometrically, is directly proportional to the B concentration in the sample (Malekani & Cresser, 1998).

Alternatively, techniques such as inductively coupled plasma optical emission spectrometry (ICP-OES) or inductively coupled plasma mass spectrometry (ICP-MS) can be employed, although these methods require more specialized instrumentation (Sah & Brown, 1997). In both cases, the low throughput and elevated cost limit the screening of large sample sets typically required in breeding programs. Furthermore, the traits related to B tolerance remains poorly understood, complicating the development of efficient screening methodologies for desirable traits such as B exclusion (Blevins & Lukaszewski, 1998).

To overcome these limitations, high-throughput phenotyping methods, which leverage proximal and remote sensing alongside algorithms for processing large, multi-dimensional datasets, are being integrated into breeding efforts to accelerate genetic gain (Araus et al., 2018; Chawade et al., 2019; Cohen & Alchanatis, 2018; Gao et al., 2022; Singh et al., 2016). Hyperspectral radiometry, in particular, has emerged as a cost-effective, non-destructive, and accurate approach for assessing key physiological traits (Sarić et al., 2022), such as nitrogen and soluble carbohydrate content in almonds (Paz-Kagan et al., 2020), chloride (Cl^-^) content in persimmons (de Paz et al., 2016) and grapevines (Sharma et al., 2024), root dry matter and starch content in cassava (Hershberger et al., 2022; Nkouaya Mbanjo et al., 2022), disease detection in wheat (Yu et al., 2018), and yield prediction in potato (Li et al., 2020).

Given the limited information on how commercially available rootstocks perform under high B levels, the interactions between B accumulation and key physiological traits, and the absence of scalable, cost-effective, and time-efficient screening methods for B toxicity, we have established two primary objectives. First, we aim to characterize B toxicity and g_s_ across varying stress levels in both commercial grapevine rootstocks and wild *Vitis* germplasm with breeding potential. Second, we seek to develop a hyperspectral-based methodology to predict B concentration, thereby accelerating breeding efforts. Our findings provide insights into how this high-throughput strategy— even with modest prediction accuracy—can outperform traditional screening techniques in breeding programs.

## Materials and Methods

### Plant materials

A selection of both commercial rootstocks and unreleased breeding materials from the University of California, Davis’s rootstock breeding program was evaluated. The commercial rootstocks included 140Ru (*V. berlandieri* × *V. rupestris*), 1103P (*V. berlandieri* × *V. rupestris*), 101-14Mgt (*V. riparia* × *V. rupestris*), Dog-Ridge (*V. champinii*), SO4 (*V. berlandieri* × *V. riparia*), Schwarzmann (*V. riparia* × *V. rupestris*), Riparia-Gloire (*V. riparia*), GRN3 ((Dog Ridge × Riparia Gloire) × *V. rufotomentosa*) × *V. champinii*), and 110R (*V. berlandieri* × *V. rupestris*). The breeding materials—T03-15 (*V. rupestris*), SAZ4 (*V. arizonica*), NM03-17-S01 (*V. treleasei*), Longii-9018 (*V. acerifolia*), Longii-9035 (*V. acerifolia*), 2014-160-003 (*V. acerifolia* × Ramsey), and 2014-160-027 (*V. acerifolia* × Ramsey)—were selected based on variation in B exclusion from previous screenings in the program.

### Experimental design

The experiment was conducted in a controlled greenhouse at the University of California, Davis. Hardwood cuttings were collected on January 8, 2024, and propagated under controlled conditions. Once rooted, the cuttings were transferred into 4-inch pots filled with fritted clay on March 15, 2024. Five B treatments were applied at concentrations of 0.5, 1, 2, 4, and 8 ppm. The experiment included six plants per genotype per treatment. B was applied through daily excess irrigation—ensuring drainage from the bottom of each pot—while all other nutrients were maintained consistently. Treatments commenced on May 7 and concluded 29 days later, on June 6, 2024, after which leaves were harvested for B analysis.

### Hyperspectral reflectance measurements

Leaf hyperspectral reflectance was measured using a CI-710s Leaf Spectrometer (CID Bio-Science, Camas, WA, USA). Measurements were taken weekly between 9:00 AM and 12:00 PM on the fourth leaf from the bottom. The raw data were preprocessed to enhance the signal-to-noise ratio as described by Hershberger et al. (2021) and then interpolated onto a uniform wavelength grid from R_380_ to R_1100_ nm (at 1-nm increments) using linear interpolation.

Four vegetation indices were computed from hyperspectral data collected on June 5, 2024, as in Roberts et al. (2018). The Normalized Difference Vegetation Index (NDVI) was calculated as (NIR − red) / (NIR + red), where red reflectance was averaged over wavelengths R_640_–R_670_ nm, and NIR reflectance was averaged over R_850_–R_880_ nm. The Photochemical Reflectance Index (PRI) was calculated as (R_531_ − R_570_) / (R_531_ + R_570_), while the Structure Insensitive Pigment Index (SIPI) was computed as (R_800_ − R_445_) / (R_800_ − R_680_). The Chlorophyll Index (CI) was determined using the formula (NIR /

RedEdge) − 1, where RedEdge reflectance was averaged over R_700_–R_720_ nm, and NIR was averaged same as for the NDVI. For each genotype–treatment combination, measurements were taken from six plants to ensure reliable data collection.

### Boron analysis

At the end of the experiment (29 days after B treatments started), all leaves were collected, oven-dried, and sent to the University of California Davis’s analytical lab. Dry samples were ground to a fine powder, and 400 mg of each sample underwent nitric acid/hydrogen peroxide microwave digestion. The digests were then analyzed by Inductively Coupled Plasma Atomic Emission Spectrometry (ICP-AES) to determine B content.

### Physiological data

g_s_ and photosystem II efficiency (ΦPSII) were measured using the LI-600 Porometer/Fluorometer (LI-COR, Lincoln, NE, USA) at the end of the experiment. For these measurements, a young, fully expanded, sun-exposed leaf was selected from each plant, with data collected on sunny days between 10:00 AM and 12:00 PM. Physiological measurements were collected only for the 0.5, 4, and 8 ppm B treatments to accommodate the high number of genotype-treatment combinations and to restrict the measurement period, thereby minimizing environmental variability.

### Statistical analysis and modeling

The spectral data were merged with the reference lab B measurements to create a comprehensive dataset. Data cleaning involved removing extreme outliers identified via principal component analysis (PCA) on the reflectance profiles. A classification threshold of 150 ppm B was used to differentiate B excluders from non-excluders, as concentrations above this level are considered toxic to grapevines (Ibacache et al., 2020). Thirteen different spectral transformation methods were applied using the ‘waves’ package (Hershberger et al., 2021). The transformations included: standard normal variate (SNV); standard normal variate and first derivative (SNVD1); standard normal variate and second derivative (SNVD2); first derivative (D1); second derivative (D2); Savitzky-Golay with window size = 11 (SG); standard normal variate and Savitzky-Golay (SNVSG); gap segment derivative with window size = 11 (SGD1); Savitzky-Golay with window size = 5 and first derivative (SG.D1W5); Savitzky-Golay with window size = 11 and first derivative (SG.D1W11); Savitzky-Golay with window size = 5 and second derivative (SG.D2W5); Savitzky-Golay with window size = 11 and second derivative (SG.D2W11).

To predict B concentrations, three different wavelength selection strategies were tested. First, prediction models were built using the full spectral range from R_380_ to R_1100_ nm. Next, highly predictive wavelengths were identified using ANOVA, with treatments and B content as the explanatory and response variables, respectively (Fig. S1), and variable importance projection (VIP) based on a partial least squares regression (PLS-R; Fig. S2). These three feature selection approaches were then evaluated using PLS-R and random forest regression (RF-R) for quantitative prediction of B, as well as partial least squares discriminant analysis (PLS-DA) and random forest classification (RF-C) for predicting B exclusion categories based on the 150 ppm threshold.

In total, 156 modeling combinations were tested, representing 13 spectral transformations, 3 feature selection strategies, and 4 prediction models. For each combination, the dataset was repeatedly partitioned into training (70%) and testing (30%) subsets, with model performance—measured as R^2^ for regression or accuracy for classification—averaged over 1000 iterations.

Using the AlphaSimR package (Gaynor et al., 2021), we simulated responses to selection across a range of intensities and accuracies to evaluate our novel B-prediction method against traditional screening approaches. In these simulations, we assumed a genetic standard deviation of 1 and evaluated heritability levels of 0.3, 0.5, 0.7, and 0.9 across population sizes of 100, 250, 500, and 1000 individuals. Selection intensity was varied by altering the number of individuals selected (ranging from 10 to 50), with intensity calculated as the ratio of the standard normal density at the truncation point to the selection proportion, scaled by the correlation between phenotype and breeding value. All modeling, statistical analyses, and data visualizations were conducted using R (version 4.4.1; R Core Team, 2024).

## Results

### Variability in B accumulation across grapevine rootstocks

The examined rootstocks and breeding materials exhibited large variability in shoot B accumulation. For instance, the leaf B content in T03-15 (*V. rupestris*) was 4.7 times higher than that in Longii-9035, which had the lowest levels. Across all genotypes, plants treated with 8 ppm B consistently showed higher leaf B content than those treated with 0.5 ppm (Fig. 1A). In half of the genotypes—namely, T03-15, 140Ru, 101-14Mgt, SAZ4, Riparia-Gloire, Longii-9018, 110R, and Longii-9035—the leaf B content at 4 ppm was also significantly higher than at 0.5 ppm (one-way ANOVAs, P < 0.05).

**Figure 1.**
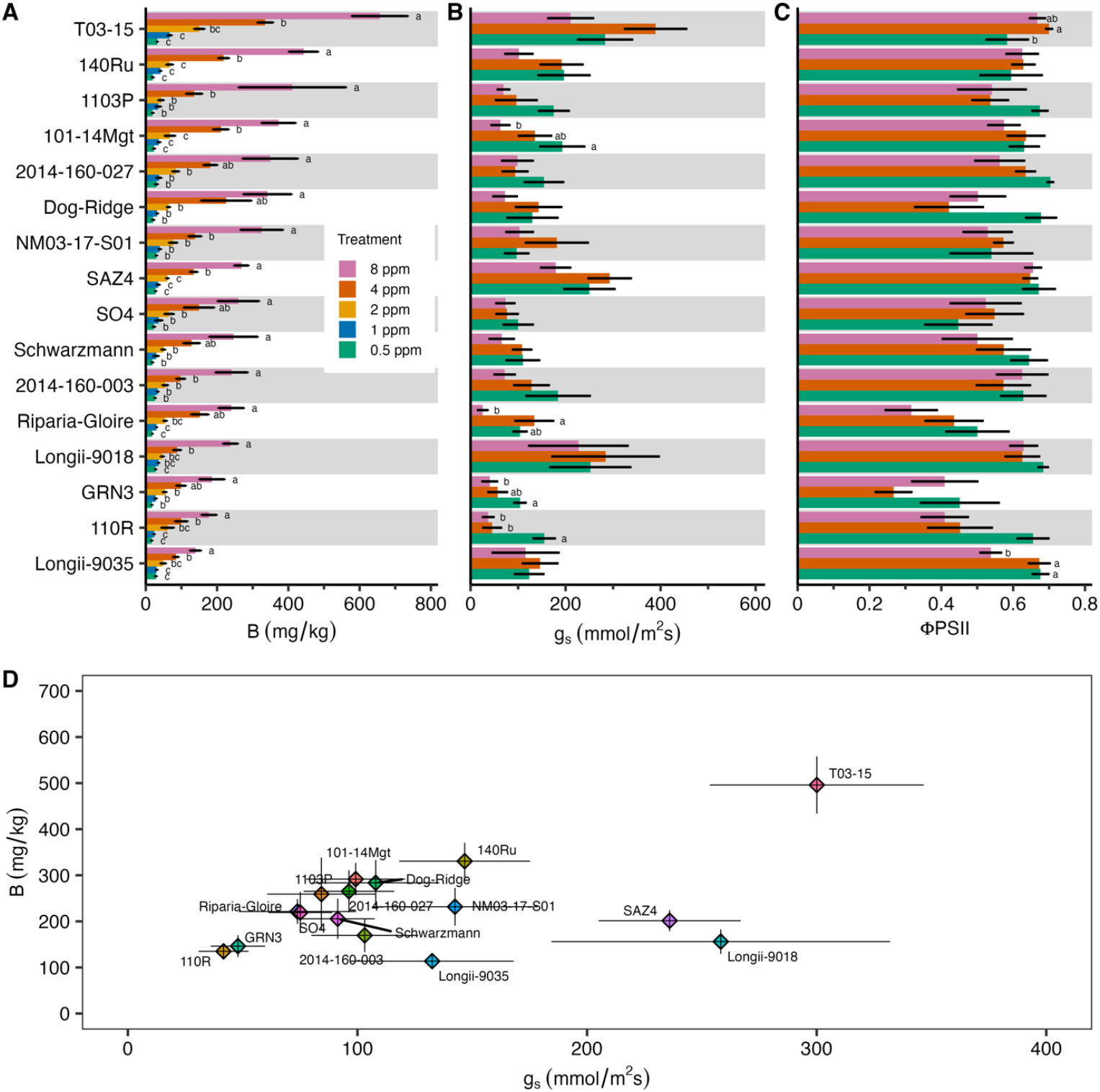
Leaf boron (B) content, stomatal conductance (g_s_), and photosystem II efficiency (ΦPSII). (A) Leaf B content, (B) g_s_, and (C) ΦPSII, ordered from highest (top) to lowest (bottom) B in the 8 ppm treatment. g_s_ and ΦPSII were only collected in the 0.5, 4 and 8 ppm treatments. Bars represent the mean ± standard error. (D) B vs. g_s_ for the 4 and 8 ppm treatments; horizontal and vertical bars represent the mean ± standard error for each variable. Different letters denote significant differences among treatments within each genotype (one-way ANOVA followed by Tuckey HSD, P < 0.05). If no letters appear for a given genotype, there were no significant differences among treatments for that genotype.

Notably, T03-15 displayed significantly higher B levels (one-way ANOVAs, P < 0.05) than all other genotypes in the 1 and 2 ppm treatments and maintained the highest B content in the 4 and 8 ppm treatments (significantly higher than most of the other genotypes; Fig. S3).

### Physiological responses and relationship with leaf B content supplementary

Overall, there was no correlation between the g_s_ and B (R^2^ < 0.01, P = 0.4). However, three genotypes, 101-14Mgt, GRN3, and 110R (commercial rootstocks, exhibited a significant reduction in g_s_ between the 8 ppm and 0.5 ppm treatments (one-way ANOVA, P < 0.05; Fig. 1B). In contrast, Longii-9035 and Longii-9018 maintained relatively consistent g_s_ across treatments. There was no correlation between ΦPSII and B (R^2^ = 0.01, P = 0.06); while some genotypes experienced a slight reduction in the 8 ppm treatment compared to 0.5 ppm, others showed no change—or even a modest increase (Fig. 1C). Notably, Longii-9035 was the only genotype that demonstrated a significant decline in ΦPSII in the 8 ppm treatment relative to the 0.5 ppm treatment (one-way ANOVA, P < 0.05).

When comparing leaf B content against g_s_, most genotypes clustered together, with several genotypes either over- or under-performing relative to the cluster (Fig. 1D). T03-15, for example, had both the highest leaf B and g_s_, whereas Longii-9035 exhibited the lowest leaf B and an intermediate g_s_. Additionally, GRN3 and 110R formed a distinct cluster with relatively low values for both parameters, while SAZ4 and Longii-9018 clustered together with low leaf B yet maintained high g_s_.

### Analysis of vegetation indices in response to B

Leaf NDVI, which serves as an estimation of leaf greenness, and PRI, which reflects changes in photosynthetic light-use efficiency, decreased in the 1, 2, 4 and 8 ppm treatments compared to the 0.5 ppm treatment (one-way ANOVA, P < 0.05; Fig. 2E, F). The SIPI, which provides insights on the balance between carotenoids and chlorophyll, decreased in the 2 and 4 ppm treatments but not in the 8 ppm treatment, compared to the 0.5 ppm treatment (Fig. 2G). The CI decreased in the 1, 2, 4, 8 ppm treatment compared to the 0.5 ppm treatment, however, it was higher in the 1 and 8 ppm, compared to the 2 and 4 ppm treatments (Fig. 2H). The genotypes had different NDVI, PRI, SIPI and CI responses to the B treatments. While most genotypes had reduced NDVI in the 8 ppm and/or 4 ppm treatments compared to the 0.5 ppm treatment, Longii-9035 had similar values of NDVI in all the treatments (Fig. 2A). Longii-9035 also had similar or higher values of PRI, SIPI and CI in the 8 ppm treatment compared to the 0.5 ppm treatment. Leaf B concentration did not correlate with any of the four vegetative indices (NDVI, PRI, CI, and SIPI; Fig. S4)

**Figure 2.**
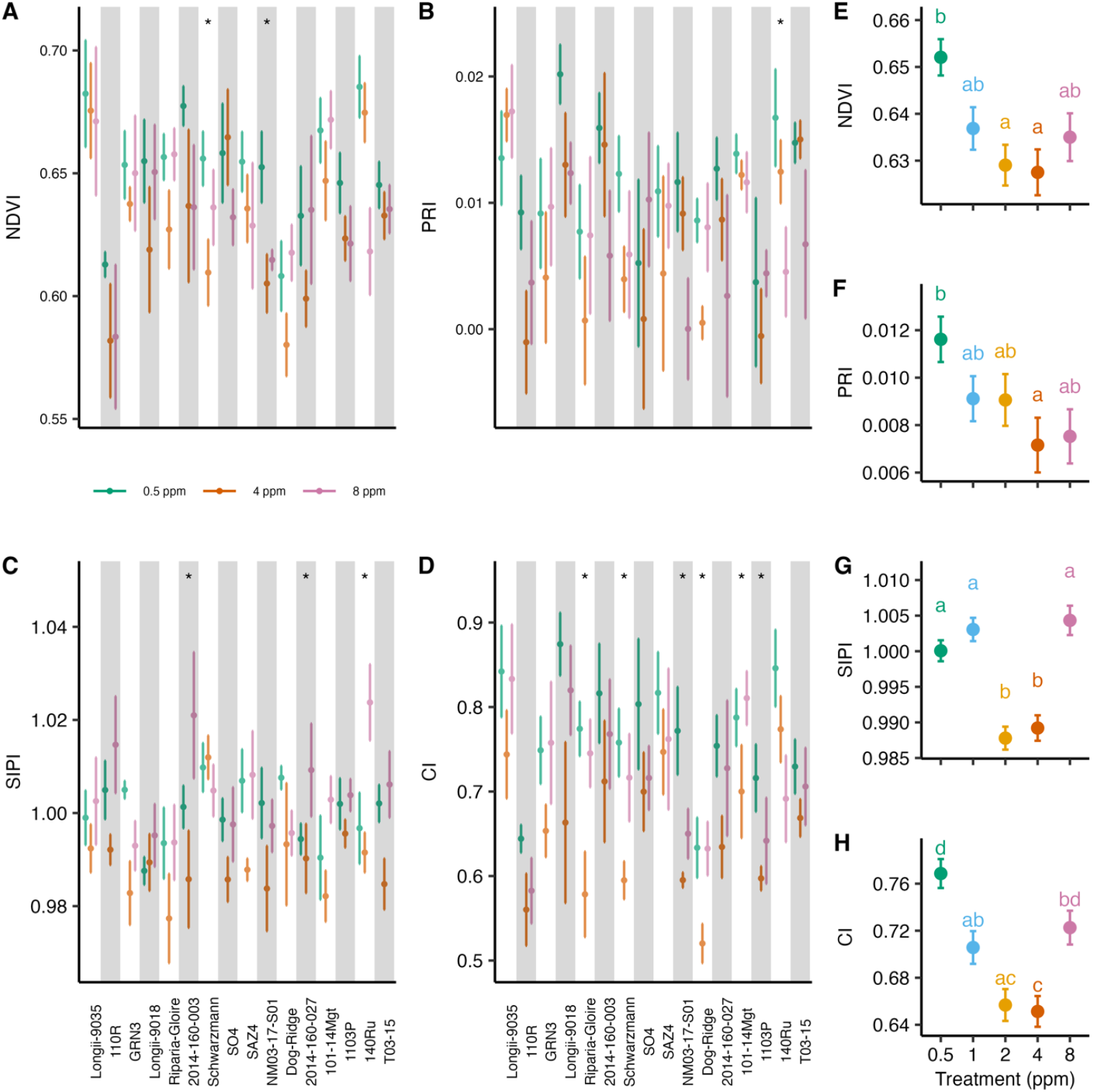
Analysis of vegetation indices derived from hyperspectral measurements. (A) Normalized Difference Vegetation Index (NDVI). (B) Photochemical Reflectance Index (PRI). (C) Structure Insensitive Pigment Index (SIPI). (D) Chlorophyll Index (CI). (E) NDVI by treatment. (F) PRI by treatment. (G) SIPI by treatment. (H) CI by treatment. Measurements were taken on June 5, 2024. Points represent the mean index values with corresponding standard errors. Different letters denote significant differences among treatments (one-way ANOVA followed by Tuckey HSD, P < 0.05). Asterisks denote significant differences among treatments within each genotype (one-way ANOVA, P < 0.05).

### Hyperspectral data processing, wavelength selection, and model development

Reflectance across most of the visible (VIS) spectrum (∼R_500_–R_700_ nm) increased with B levels, showing the lowest reflectance in the 0.5 ppm treatment and the highest in the 8 ppm treatment. However, this pattern did not hold in NIR region (∼R_750_-R_1100_ nm), where the lowest (0.5 ppm) and highest (8 ppm) treatments exhibited higher reflectance than the intermediate treatments (1, 2 and 4 ppm; Fig. 3A). One-way ANOVAs performed separately for each wavelength revealed significant differences among the B treatments for most of the examined spectral range (∼R_475_–R_520_ nm and above R_600_ nm; P < 0.05; Fig. 3B). Examination of VIP scores calculated from a PLS-R model revealed a different pattern, where VIP scores greater than 1, indicative of high predictive relevance, occurred mostly at the edges of the spectrum, including a notable peak at the edge of the VIS region (∼R_630_–R_760_ nm; Fig. 3C).

**Figure 3.**
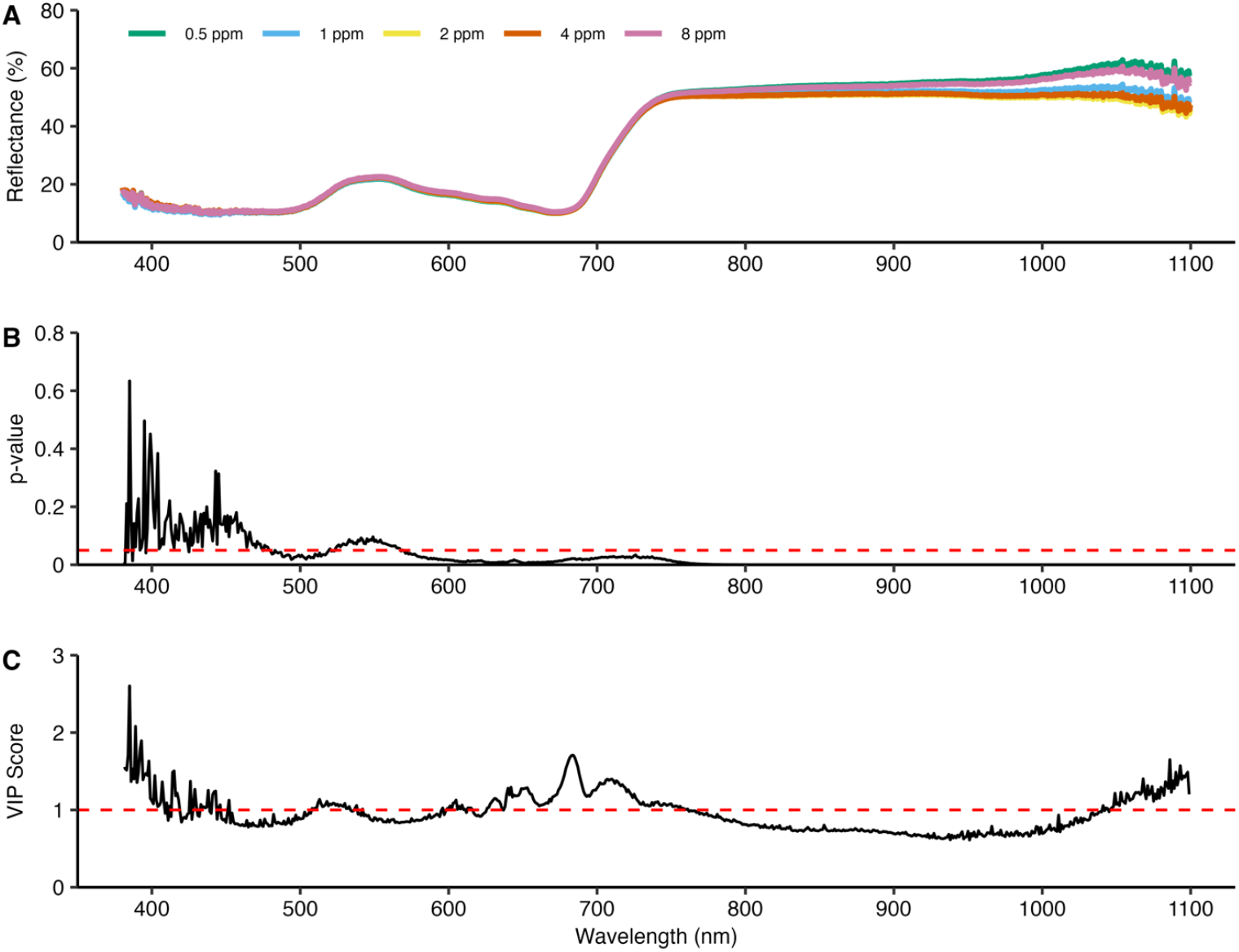
Raw reflectance and results of the ANOVA and variable importance projection (VIP) based selection criteria. (A) Raw reflectance for each of the B treatments. Each line is the mean of all the plants from one treatment. (B) P-values from one-way ANOVAs between the B treatment within each wavelength. Wavelengths with P < 0.05 (below the red dotted line) were selected for prediction. (C) VIP scores calculated from a preliminary PLS model between the raw reflectance data and the measured B. Wavelengths with VIP > 1 (above the red dotted line) were selected for prediction.

### Evaluation of regression and classification models for prediction of leaf B

We observed clear performance differences between models predicting quantitative B concentrations (PLS-R and RF-R) and those classifying B exclusion categories (PLS-DA and RF-C). PLS-R and RF-R exhibited consistently lower accuracy (∼0.2 to 0.4) regardless of the wavelength selection strategy (using all wavelengths, or selecting wavelengths based on ANOVA or VIP) and the spectral transformation method applied (Fig. 4 A and B). In contrast, models focused on categorical prediction, PLS-DA and RF-C, achieved substantially higher accuracy (∼0.7 to 0.9), suggesting that spectral data are more effective at distinguishing B-excluding genotypes than at precisely quantifying B concentrations (Fig. 4 C and D).

**Figure 4.**
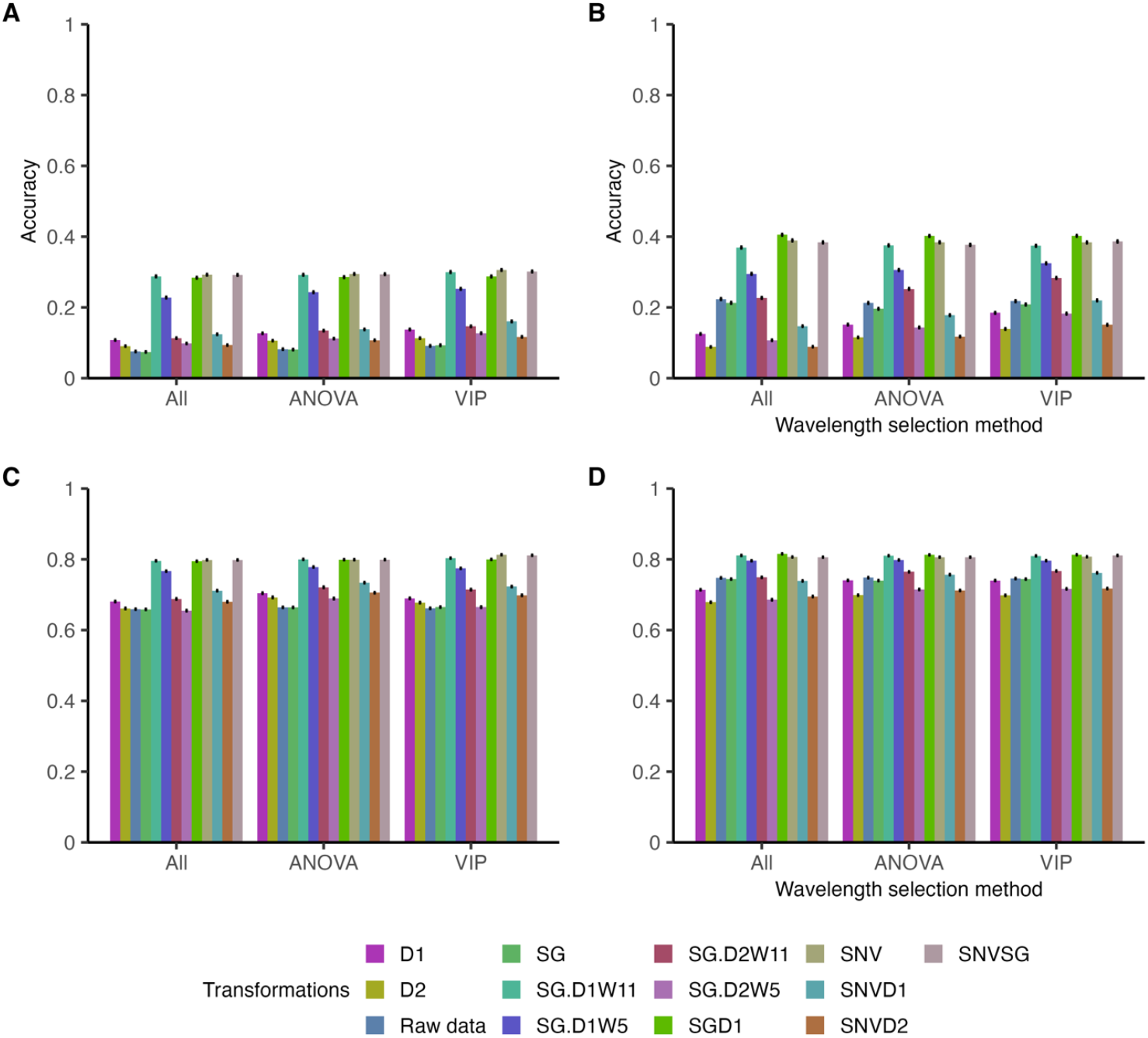
Performance accuracies of boron (B) prediction models for different wavelength selection strategies and transformations methods. (A) Partial Least Squares Regression (PLSR). (B) Random Forest regression model (RF-R). (C) Partial Least Squares Discriminant Analysis (PLS-DA). (D) RF classification model (RF-C). Models are compared under three wavelength-selection criteria: All: using all available wavelengths; ANOVA: using only wavelengths with P < 0.05 from an ANOVA between treatments; VIP: using only wavelengths with VIP score > 1 from a PLSR model. For each combination, different colors represent different reflectance transformations. Each bar shows the mean performance accuracy over 1000 iterations of each model. Error bars indicate the standard errors of the mean. For PLSR and RF regression, performance is reported as R2; for the PLS-DA and RF classification models, accuracy corresponds to classification performance. The number of trees for both RF models was 500. Measurements were taken after 29 days of treatment.

Across all models, the choice of wavelength selection method had minimal impact on performance, with accuracy remaining relatively stable whether all wavelengths were used, or features were selected via ANOVA or VIP. Similarly, the spectral transformation method did not produce large deviations in accuracy, though some transformations, such as SG.D1W11, SG.D1W5, SNV, and SNVSG, tended to yield slightly better performance in PLS-DA and RF-C.

To assess the potential for early B prediction and classification, we applied similar modeling approaches to reflectance data collected eight days after treatments began (Fig. 5). These models followed the same methodology as those used for the final hyperspectral measurements (Fig. 4). As observed with the final measurements, regression models had lower accuracy than classification models. Among the regression approaches, PLS-R showed the poorest performance, with a maximum R^2^ of 0.21 (Fig. 5A), whereas RF-R performed slightly better, reaching a maximum R^2^ of 0.34 (Fig. 5B). Compared to the final hyperspectral measurement, PLS-R had a similar range of low accuracy, while RF-R had a slightly lower average R^2^ in the early measurement (0.11) than in the final measurement (0.26, Fig. 4C, 4C). However, the best-performing RF-R model using early measurement data (SNV, all wavelengths, R^2^ = 0.34) showed comparable accuracy to the best RF-R model from the final measurement (SGD1, all wavelengths, R^2^ = 0.40; Fig. 5C).

**Figure 5.**
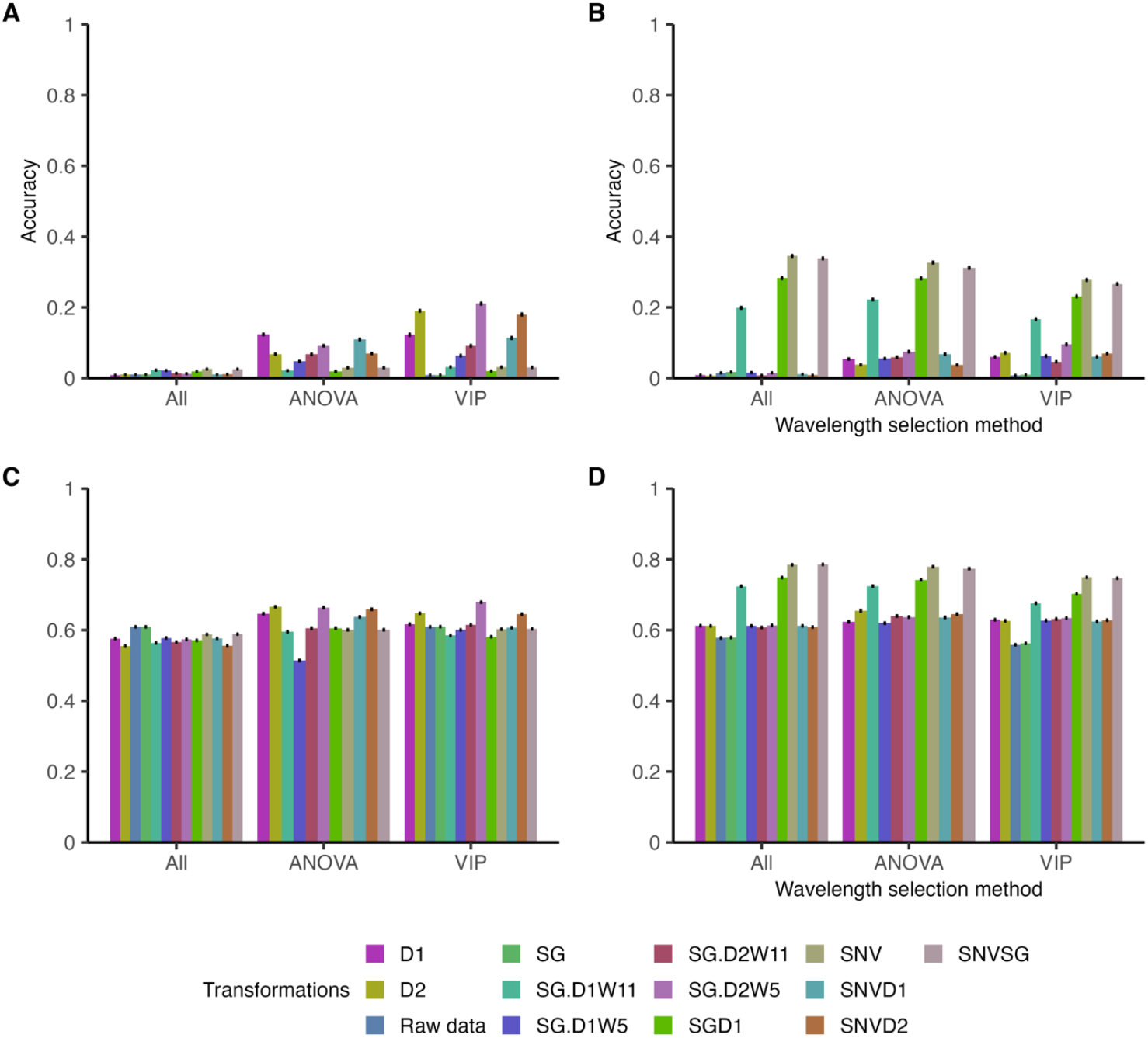
Performance accuracies of boron (B) prediction models after 8 days of B treatment. (A) Partial Least Squares Regression (PLS-R). (B) Random Forest regression model (RF-R). (C) Partial Least Squares Discriminant Analysis (PLS-DA). (D) RF classification model (RF-C). Models’ methodologies are explained in Fig. 4 and in the materials and methods. For PLS-R and RF-R, performance is reported as R^2^; for the PLS-DA and RF-C models, accuracy corresponds to classification performance. Measurements were taken after 8 days of treatments, on May 15, 2024.

For classification models, PLS-DA performed worse with early measurement data, achieving a maximum accuracy of 0.68 (Fig. 5B), compared to 0.81 in the final measurement (Fig. 4B). RF-C also showed a drop in accuracy, with an early measurement average of 0.66 (Fig. 5D) compared to 0.76 in the final measurement (Fig. 4D). However, SNVSG and SNV transformations maintained relatively stable performance, with early measurement accuracies of 0.79 and 0.78, respectively, compared to 0.81 for both transformations in the final measurement. These results indicate that while early spectral measurements can capture some variation in B content, classification models remain more reliable than regression models for early B prediction.

## Discussion

The genotypes examined exhibited large variation in leaf B exclusion, which allows for breeding new B-tolerant rootstocks. Contrary to expectations, there was no correlation between g_s_ and leaf B content (Fig. S5). Since B is primarily transported to the shoots via the xylem stream (Pereira et al., 2021), we anticipated that higher g_s_ would lead to increased B translocation and accumulation in the leaves. However, T03-15, which had the highest leaf B content, also exhibited the highest g_s_, while GRN3 and 110R, which had relatively low B content, showed the lowest g_s_.

Among the genotypes with low leaf B, SAZ4 and Longii-9018 maintained relatively high g_s_, suggesting that their lower B content may be due to active B exclusion mechanisms rather than reduced transpiration. These exclusion mechanisms could involve reduced B uptake from the soil or enhanced B removal from the shoot (Brown & Hu, 1996; Miwa et al., 2007; Wakuta et al., 2016), and might involve aquaporins (Mosa et al., 2016). B and Cl^-^ stresses share similarities, as both ions function as micronutrients but become toxic at high concentrations (Munns & Tester, 2008). However, B and Cl^-^ differ in their toxic threshold concentrations and mobility within the plant, as B is generally considered less mobile than Cl^-^ (Brown & Shelp, 1997; White & Broadley, 2001).

Previous studies have identified a combined response to B and Cl^-^ stress in grapevines (Ben-Gal et al., 2008; Downton & Hawker, 1980), and *Prunus* rootstocks (El-Motaium et al., 1994). Notably, Longii-9035 and Longii-9018, which exhibited low B accumulation, also demonstrated enhanced Cl^-^ exclusion capacities (Sharma et al., 2024). The ability of these genotypes to exclude both B and Cl^-^ under high concentrations suggests that a shared physiological mechanism may be involved. These findings underscore their potential as valuable candidates for future breeding programs aimed at developing rootstocks with combined B and Cl^-^ tolerance.

The positive correlation between increased reflectance in the VIS range and higher B levels (Fig. 3A) suggests that elevated B concentrations reduce light absorption by the leaves. This effect may be linked to decreased photosynthetic activity due to B toxicity, which can reduce chlorophyll content or increase carotenoid accumulation (Landi et al., 2013). However, this trend did not persist in the VNIR range, where the lowest (0.5 ppm) and highest (8 ppm) B treatments exhibited higher reflectance than the intermediate treatments (1, 2, and 4 ppm). One possible explanation for the 8 ppm treatment is that excessive B accumulates in the mesophyll cell wall, leading to denser cell walls that reduce light absorption in the VNIR range (Ozturk et al., 2010). Previously, high concentrations of B were found to change functional groups in rice leaves’ cell walls, such as the methine group (-CH), which affects the structural and functional roles of cellulose and pectin (Riaz et al., 2021).

Our results demonstrate that hyperspectral radiometry combined with machine learning is a promising approach for large-scale, cost-effective, and time-efficient B prediction. Classification models for B exclusion demonstrated strong potential, achieving moderate to high accuracy even within a few days of stress initiation. This offers a practical solution for screening plant material in breeding programs, especially when phenotyping is a bottleneck. Similar results were found in the classification of virus infection in grapevines using hyperspectral sensing (Wang et al., 2023). In contrast, quantitative B prediction was less reliable, with regression models yielding low to moderate accuracy. These findings align with previous research showing that micronutrient concentrations, including B, are more challenging to predict from hyperspectral data than macronutrients (Grieco et al., 2022; Pandey et al., 2017). In fact, Pandey et al. (2017) identified B as one of the least predictable elements. Similarly, Sharma et al. (2024) found that classification models were more effective than regression models for predicting Cl^-^ exclusion in grapevines using hyperspectral data, suggesting that categorical approaches may be more robust for ion exclusion studies.

The observed reductions in NDVI and PRI with increasing B concentration in the irrigation solution suggest a decline in photosynthetic efficiency, possibly due to chlorophyll loss (Gitelson & Merzlyak, 1997) or reduced light-use efficiency in photosynthesis (Gamon et al., 1992). The reduction in CI further supports the idea that NDVI declines were driven by reduced chlorophyll content (Gitelson & Merzlyak, 1994). Interestingly, SIPI followed a different trend, decreasing at 2 and 4 ppm but not at 8 ppm. This distinct response may indicate increased carotenoid accumulation in the leaves, as suggested by Penuelas et al. (1995). The differences in SIPI and CI patterns between the 2 and 4 ppm treatments versus the 8 ppm treatment suggest that at high B concentrations, carotenoid accumulation may serve as a protective mechanism against B-induced stress, a phenomenon observed under other stress conditions as well (Uarrota et al., 2018). These findings confirm that B stress negatively impacts photosynthesis and suggest that carotenoid accumulation may be a tolerance strategy to mitigate B toxicity. This insight could help identify genotypes with improved B stress resilience, informing future breeding efforts for B-tolerant rootstocks.

Moderate accuracies (e.g., 0.5–0.6) may limit the appeal of radiometry-based methods for certain breeding applications. However, in a breeding context—particularly when considering response to selection—lower accuracies can be offset by larger population sizes and higher selection intensities. To explore this potential, we simulated how different population sizes, selection intensities, and heritabilities affect the response to selection (R; Fig. 6). The results follow expected patterns: R increases with rising heritability and greater selection intensity. By removing the phenotyping bottleneck—for instance, through cost-effective, high-throughput spectroscopy—breeders can expand their screening capacity from 100 to 500 individuals. Under these conditions, tripling the selection intensity from 10 to 30 individuals increases R from about 1.5 to 2, reflecting roughly a 33% improvement in genetic gain. Although these simulations are primarily illustrative and do not account for detailed cost analyses of different screening technologies, they highlight how increasing population size and adjusting selection intensity can substantially boost the response to selection.

**Figure 6.**
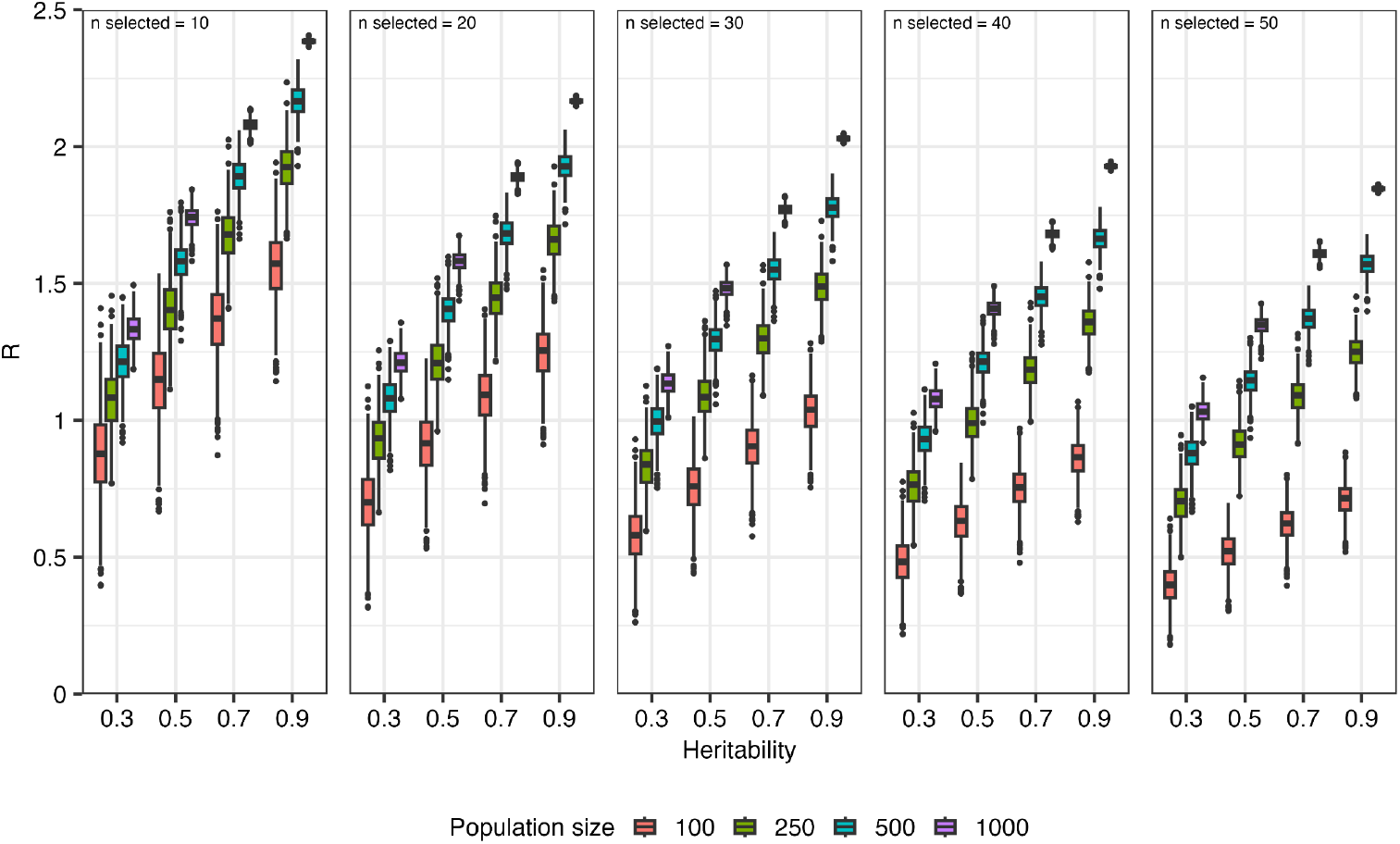
Response to selection (R) as a function of heritability, population size, and selection intensity. Boxplots display the distribution of the simulated genetic response (R) across 4 heritability levels (0.3, 0.5, 0.7, and 0.9) for 4 population sizes (100, 250, 500, and 1000 individuals) over 1000 iterations. Each panel corresponds to a fixed number of individuals selected (n selected), with the selection intensity calculated as the ratio of the standard normal density at the truncation point to the selection proportion, multiplied by the correlation between phenotype and breeding value and a genetic standard deviation of 1. Different colors indicate the distinct population sizes.

## Conclusions

Our study highlights the substantial variation in leaf B accumulation across different *Vitis* genotypes, demonstrating clear opportunities to breed new B-tolerant rootstocks. Although we initially expected a link between g_s_ and B uptake, no such correlation emerged, suggesting that active exclusion mechanisms may play a larger role than transpiration-driven processes in controlling B accumulation. Notably, several genotypes (e.g., Longii-9018, Longii-9035, SAZ4) combined low B accumulation with robust physiological performance, and some even showed improved Cl^-^ exclusion, hinting at a shared mechanism for multi-ion tolerance. Vegetation indices further confirmed that B stress impairs photosynthesis—through chlorophyll depletion or increased carotenoid accumulation—underscoring the potential for using these non-destructive metrics to track stress responses. Our findings also show that hyperspectral radiometry integrated with machine learning can effectively classify B-excluding genotypes, offering a scalable, cost-effective, and time-efficient phenotyping approach. While regression models for quantitative B prediction remain less reliable, classification models proved sufficiently accurate for early screening—critical for breeding programs where phenotyping is often a bottleneck. Moreover, simulation results indicate that moderate predictive accuracies can still drive substantial genetic gain if population sizes, and selection intensities are increased. By removing or reducing the phenotyping bottleneck, breeders can more readily exploit genetic diversity for B tolerance, accelerating the development of robust, multi-stress-tolerant rootstocks that maintain productivity under high B and other abiotic stresses.

## Acknowledgments

The authors would like to thank Mikayla Bailey, Guillermo Garcia-Zamora, and Patrick H. Brown for their support and valuable recommendations in the execution of these experiments.

## Data Availability Statement

All the collected data have been made available in the supplementary files accompanying this manuscript.

## Supplementary Information

**Figure S1.**
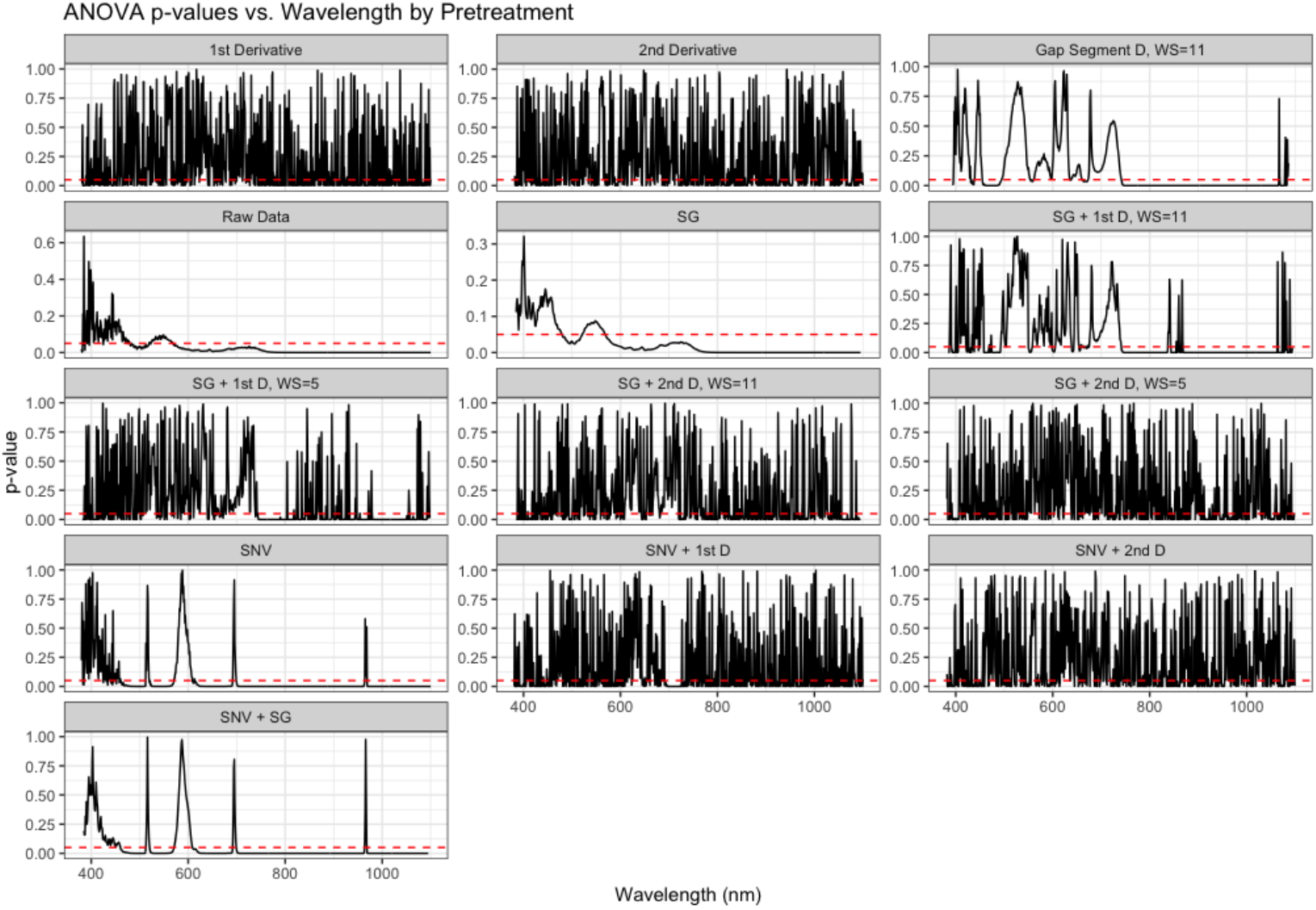
P-values obtained from one-way ANOVAs comparing spectral reflectance among different boron (B) treatments for each wavelength across the measured spectrum. Each panel represents a different spectral data transformation method performed using the R waves package. The horizontal dashed line (P = 0.05) indicates the threshold for statistical significance. Wavelengths with P-values below this threshold (highlighted regions) were considered significantly influenced by B treatments and were selected for subsequent predictive modeling analyses.

**Figure S2.**
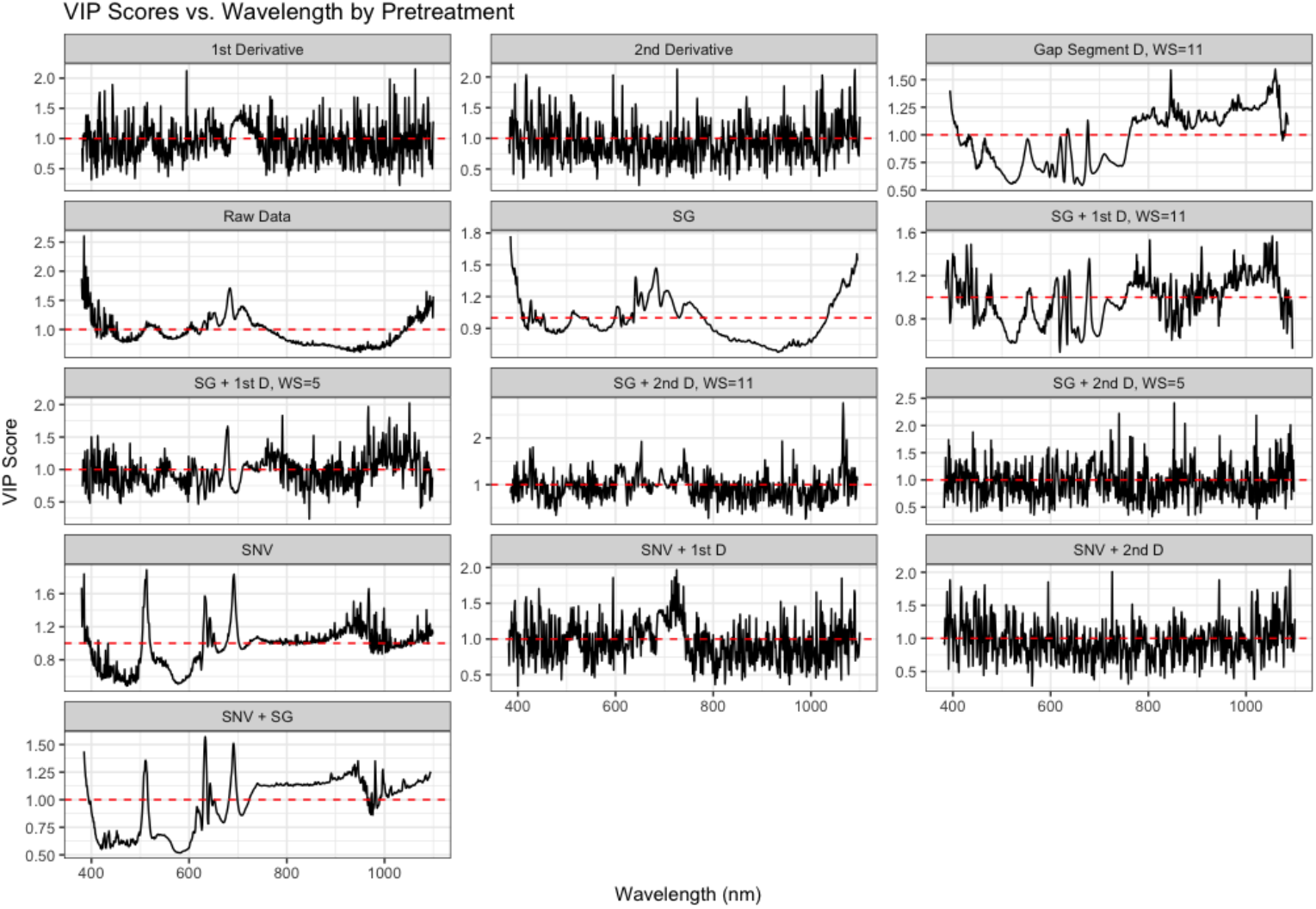
Variable importance projection (VIP) scores derived from a partial least squares regression (PLS-R) relating spectral reflectance to leaf boron (B) concentrations. Each panel represents a different spectral data transformation method performed using the R waves package. VIP scores quantify the relative contribution of each wavelength to predicting B levels. The dashed horizontal line (VIP = 1) indicates the threshold for high predictive relevance; wavelengths exceeding this threshold significantly contribute to the model’s predictive power and were selected for further analysis.

**Figure S3.**
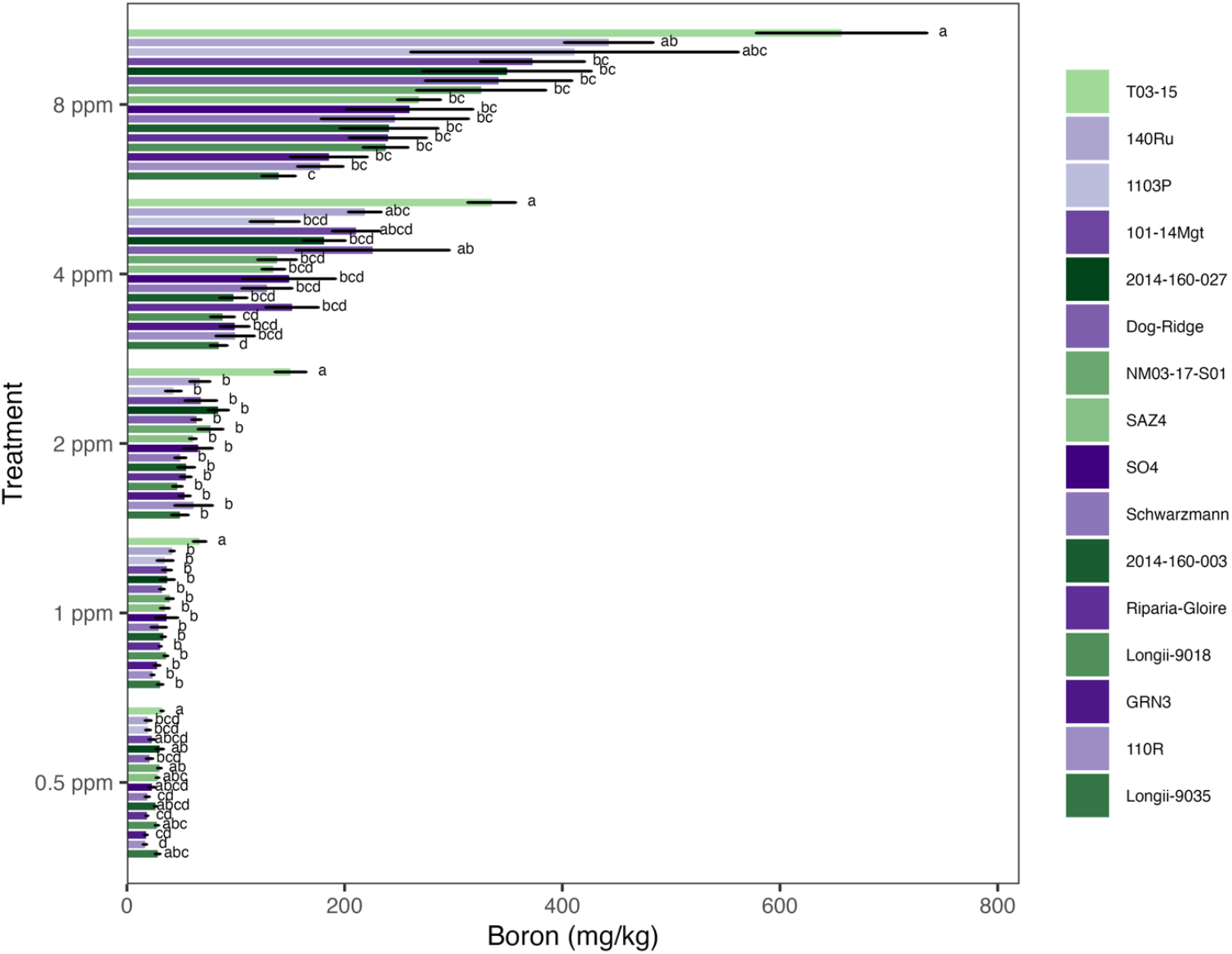
Leaf boron (B) content across genotypes. Bars represent mean ± standard error (n = 6). Different letters denote significant differences among genotypes within each B treatment (one-way ANOVA, P < 0.05). Green bars represent commercial rootstocks, while purple bars represent breeding materials.

**Figure S4.**
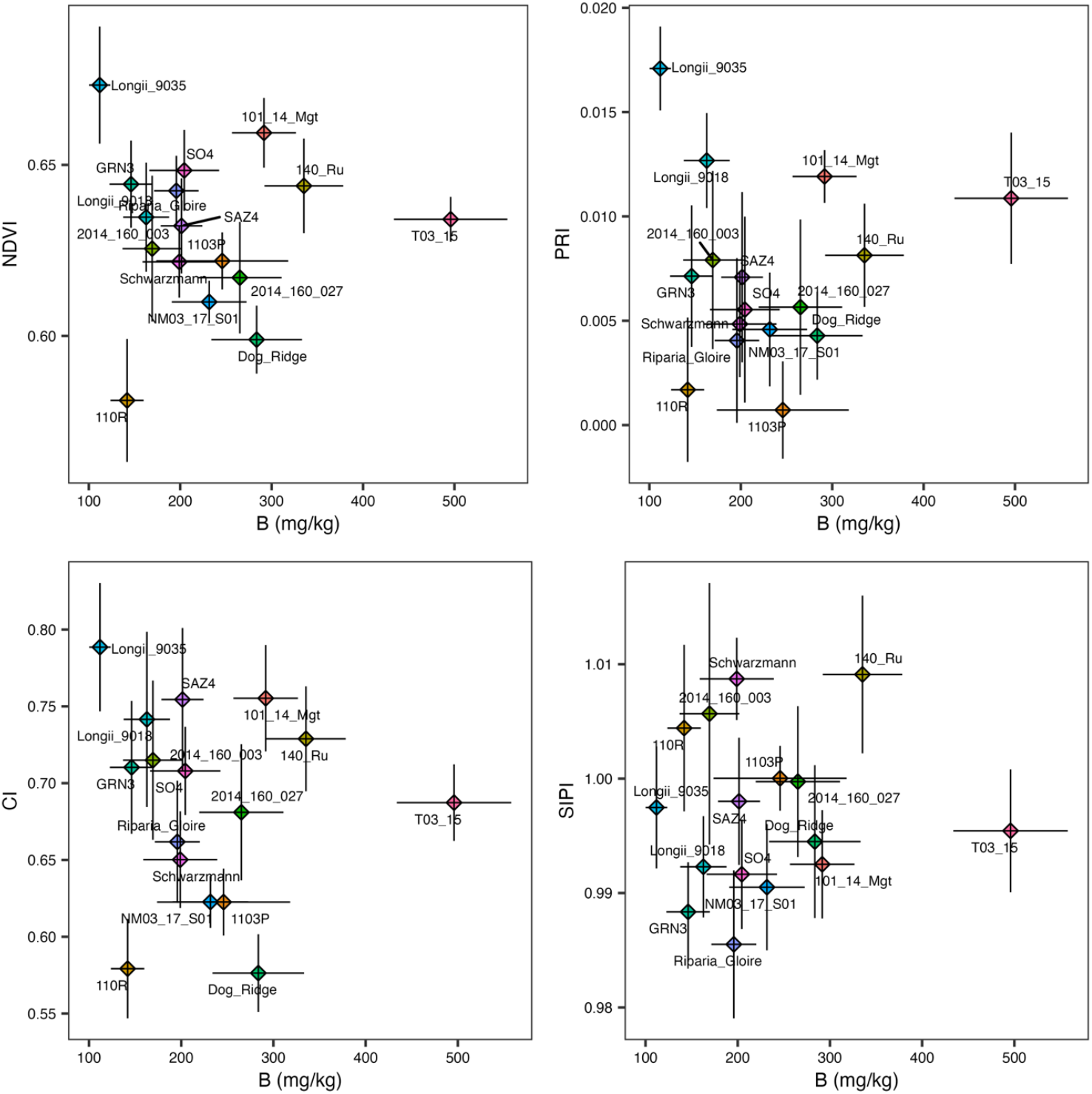
Leaf boron (B) content vs. different vegetation indices derived from hyperspectral measurements. (A) B vs. Normalized Difference Vegetation Index (NDVI). (B) B vs. Photochemical Reflectance Index (PRI). (C) B vs. Structure Insensitive Pigment Index (SIPI). (D) B vs. Chlorophyll Index (CI). Measurements were taken on June 5, 2024. Points represent the mean B and index values with corresponding standard errors. The 4 and 8 ppm treatments were used in this analysis.

**Figure S5.**
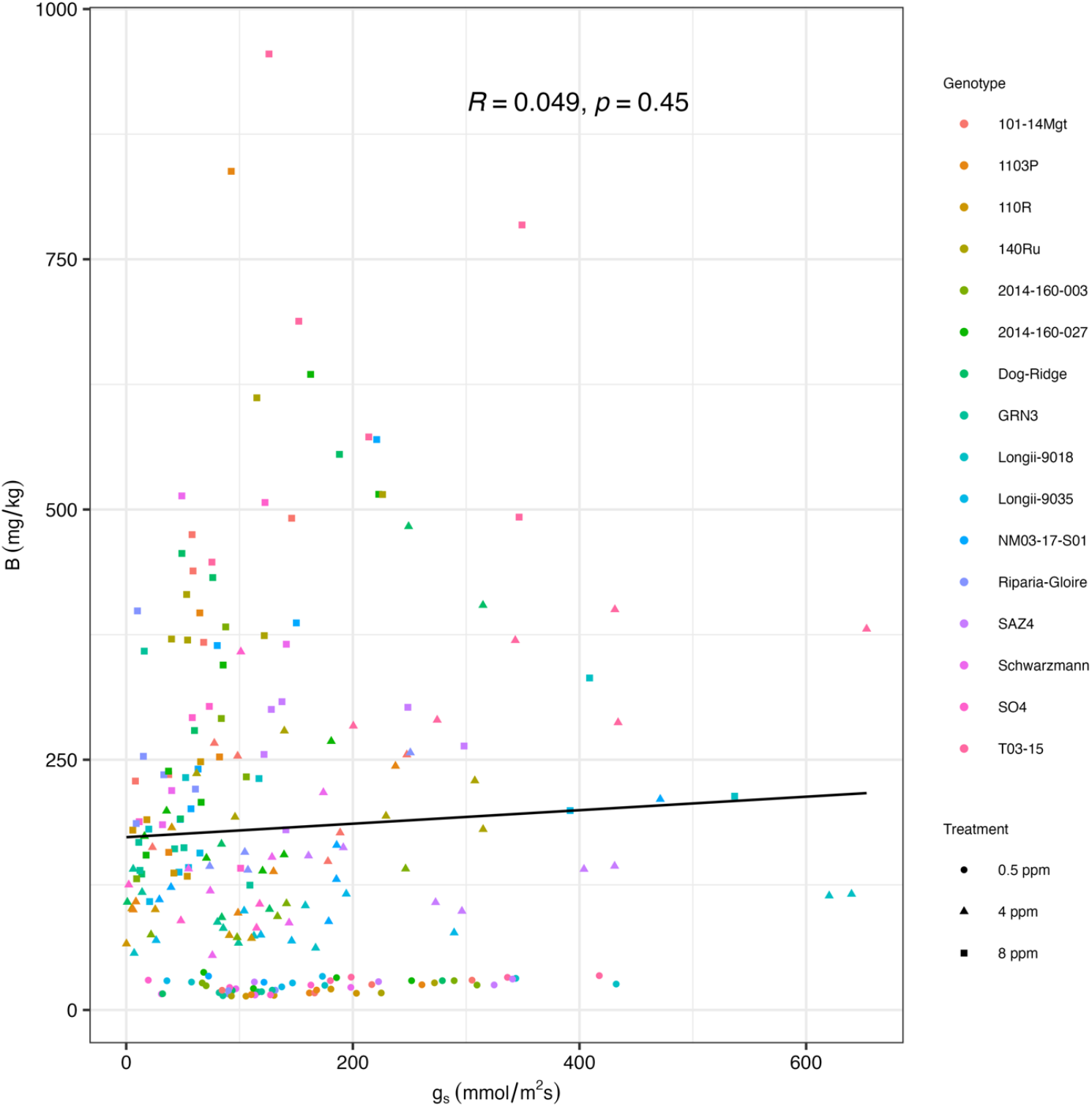
Scatter plot illustrating the relationship between leaf boron (B) concentration and stomatal conductance (g_s_). Colors indicating genotypes and shapes indicating treatments. A fitted linear regression line is shown to highlight the overall correlation between leaf B concentration and g_s_.

